# High throughput estimation of functional cell activities reveals disease mechanisms and predicts relevant clinical outcomes

**DOI:** 10.1101/076083

**Authors:** Marta R. Hidalgo, Cankut Cubuk, Alicia Amadoz, Francisco Salavert, José Carbonell-Caballero, Joaquin Dopazo

**Affiliations:** Computational Genomics Department, Centro de Investigación Príncipe Felipe (CIPF), C/ Eduardo Primo Yufera 3, Valencia, 46012, Spain; Functional Genomics Node (INB), C/ Eduardo Primo Yufera 3, Valencia, 46012, Spain; Bioinformatics in Rare Diseases (BiER), Centro de Investigación Biomédica en Red de Enfermedades Raras (CIBERER), C/ Eduardo Primo Yufera 3, Valencia, 46012, Spain

**Keywords:** Signaling pathway, disease mechanism, prognostic, survival, biomarker

## Abstract

Understanding the aspects of the cell functionality that account for disease or drug action mechanisms is a main challenge for precision medicine. Here we propose a new method that models cell signaling using biological knowledge on signal transduction. The method recodes individual gene expression values (and/or gene mutations) into accurate measurements of changes in the activity of signaling circuits, which ultimately constitute high-throughput estimations of cell functionalities caused by gene activity within the pathway. Moreover, such estimations can be obtained either at cohort-level, in case/control comparisons, or personalized for individual patients. The accuracy of the method is demonstrated in an extensive analysis involving 5640 patients from 12 different cancer types. Circuit activity measurements not only have a high diagnostic value but also can be related to relevant disease outcomes such as survival, and can be used to assess therapeutic interventions.

## Introduction

Despite most phenotypic traits (including disease and drug response) are multi-genic, the vast majority of biomarkers in use are based on unique gene alterations (expression changes, mutations, etc.) Obviously, the determination of the status of a single gene is technically easier than multiple gene measurements. However, regardless of their extensive clinical utility, single gene biomarkers frequently lack any mechanistic link to the fundamental cellular processes responsible for disease progression or therapeutic response. Such processes are better understood as pathological alterations in the normal operation of functional modules caused by different combinations of gene perturbations (mutations or gene expression changes) rather than by alterations of a unique gene [1].

Of particular interest are signaling pathways, a type of functional module known to play a key role in cancer origin and progression, as well as in other diseases. Consequently, analysis of the activity of signaling pathways should provide a more informative insight of cellular function. Actually, the recent demonstration that the activity of a pathway presents a significantly better association to bad prognostic in neuroblastoma patients than the activity of their constituent genes (among them *MICN*, the conventional biomarker) [2] constitutes an elegant confirmation of this concept. In a similar example drug sensitivity is shown to be better predicted using probabilistic signaling pathway models than directly using gene activity values [3].

However, conventional methods for pathway analysis, even the most sophisticated ones based on pathway topology, can only detect the existence of a significant level of gene activity within the pathway [4]. However, these methods ignore the obvious fact that many pathways are multifunctional and often trigger opposite functions (e.g. depending the receptor and the effector proteins involved in the transduction of the signal, the apoptosis pathway may trigger survival or cell death). Moreover, whether the level of gene activity detected by conventional methods actually triggers cell functionalities or not and, if so, what genes are the ultimate responsible for the resulting cell activity is something that must be determined *a posteriori,* usually by heuristic methods. Thus, pathway activity analysis (PAA) emerges as an alternative way of defining a new class of mechanistic biomarkers, whose activity is related to the molecular mechanisms that account for disease progression or drug response. However, capturing the aspects of the activity of the pathway that are really related to cell functionality is not trivial. This requires of an appropriate description of the elementary sub-pathways and an adequate computation of the individual contributions of gene activities to the actual activity of the sub-pathway. Different ways of computing activity scores for diverse sub-pathway definitions using gene expression values [5-8], or even gene mutations [9], have been proposed (See Table 1). However, in most of them sub-pathway definition is either unconnected, or only collaterally related, to the functional consequences of pathway activity (See Table 1).

**Table 1.** List of methods for Pathway Analysis. The first column (Method) contains the name or acronym of the method, if exists, otherwise, we refer to it as the fires author of the publication. The second column (Date) contains the publication date. The third column (code) informs on the availability of the code to run the method. The fourth column (Pathway modelled) indicates the pathway definition used in the method. The fifth column (Entity modelled) is the entity, within the pathway, used in the method (“subpath identification” methods obtain candidate sub-pathways usually by differential expression of its constituent genes, “signal quantification” methods provide, in addition, a quantification of the activation status of the sub-pathway). The sixth column (input) indicates the data type that inputs the method (MA: Expression Microarray; CNV: copy number variation; NA: not available). The seventh column (output) describes the results provided by the method. Some provide only a score (p-value, DE: differential expression matrix; PF: perturbation factor) for the whole pathway and other also provide scores for sub-pathways, that can be defined within the pathways in many different ways. The eight column (Comparison) indicates the type of comparison the method can deal with. It can be either a conventional two conditions (typically case/control) comparison or it can allow obtaining personalized results per individual. And the ninth column (Loops) indicates whether the method can handle loop structures in the topology of the sub-pathway analyzed or not.

| Method | Date | Code | Pathway modelled | Entity modelled | Input | Output | Comparison | Loops |
| --- | --- | --- | --- | --- | --- | --- | --- | --- |
| MinePath[52] | 2015 | Web application<br>http://minepath.org/ | KEGG pathways | Subpath identification | MA | p-value per pathway<br>p-value per subpathway<br>binary value per sample<br>graphical visualization | Two conditions | NA |
| Qin et al.[53] | 2015 | NA<sup>b</sup> | 12 cancer-related KEGG pathways | signal quantification | Mutations<br>CNVs<br>Cancer drugs | Pathway activity | Personalized | yes |
| subSPIA[32] | 2015 | R code | KEGG pathways | signal quantification | MA<br>RNAseq (via SPIA in ToPaSeq) | p-value of DE per subpathway<br>p-value of PF per subpathway<br>global p-value (DE+PF) | Two conditions | no |
| Pathome[54] | 2014 | NA | KEGG pathways | signal quantification | MA<br>RNAseq | p-value per subpathway | Two conditions | NA |
| Pepe et al.[55] | 2014 | R code | KEGG pathways | subpath identification | MA | p-value per subpathway | Two conditions | NA |
| ToPaSeq[41] | 2014 | R package | graphite gene-gene | integrates other | MA | Depends on the method | Two | Depends |
|  |  |  | networks<br>user's pathways | methods:<br>TopologyGSA<br>DEGraph<br>Clipper<br>SPIA<br>TAPPA<br>PRS<br>PWEA | RNAseq |  | conditions | on the<br>method |
| DEAP[38] | 2013 | python code | user defined pathway<br>structure | signal<br>quantification | MA<br>RNAseq | Score and p-value per<br>pathway<br>subgraph with the<br>maximum absolute score | Two<br>conditions | yes |
| CliPPER[5] | 2013 | R package<br>ToPASEq R package | graphite gene-gene<br>networks<br>cliques<br>user's pathways (via<br>ToPASEq) | subpath<br>identification | MA<br>RNAseq | p-value at pathway level<br>Most affected subgraph<br>per pathway<br>Gene-level statistics for<br>DE of genes | Two<br>conditions | no |
| GraphiteWeb[56] | 2013 | Web application:<br>http://graphiteweb.bio.unipd.it/<br>R package | KEGG pathways<br>Reactome pathways | integrates other<br>methods:<br>Hypergeometric<br>test<br>Global Test<br>GSEA<br>SPIA<br>CliPPER | MA<br>RNAseq | Significant pathways<br>Visualization of the<br>pathways with nodes<br>coloured according to<br>their contribution to the<br>analysis | Two<br>conditions | no |
| TEAK[57] | 2013 | Code @ Google (Windows<br>and Mac) | KEGG pathways | metabolism-<br>oriented<br>subpathway<br>identification | MA | Ranked subpathways | Two<br>conditions | no |
| PRS[40] | 2012 | ToPASEq R package | graphite gene-gene<br>networks (ToPASEq)<br>user's pathways (via<br>ToPASEq) | pathway<br>identification | MA<br>RNAseq | p-value per pathway<br>gene-level statistics for<br>DE of genes | Two<br>conditions | yes |
| DEGraph[6] | 2012 | R package<br>ToPASeq R package | subgraphs of a large graph (branch-and-bound-like approach)<br>graphite gene-gene networks (ToPASeq)<br>user's pathways (via ToPASeq) | subpath identification | MA<br>RNAseq | p-value of DE per subpathway<br>p-value per pathway<br>Gene-level statistics for DE of genes | Two conditions | no |
| Rivera et al.[58] | 2012 | NA | NetPathpathways | subpath identification | MA | p-value of most perturbed subpathway | Two conditions | NA |
| Chen et al.[59] | 2011 | NA | KEGG pathways | subpath identification | MA | p-value per subpathway<br>p-value of key genes | Two conditions | NA |
| PWEA[30] | 2010 | ToPASeq R package | Complete pathways (KEGG)<br>graphite gene-gene networks (ToPASeq)<br>user's pathways (via ToPASeq) | pathway identification | MA<br>RNAseq | p-value of DE per pathway<br>Gene-level statistics for DE of genes | Two conditions | no |
| TopologyGSA[31] | 2010 | ToPASeq R package | Complete pathways (KEGG)<br>Cliques<br>graphite gene-gene networks (ToPASeq)<br>user's pathways (via ToPASeq) | subpath identification | MA<br>RNAseq | p-value of DE per pathway<br>Gene-level statistics for DE of genes | Two conditions | no |
| DEGAS[60] | 2010 | Java (Windows) | KEGG pathways<br>PPIs network | novel subpath identification | MA | A subpathway per pathway | Two conditions | NA |
| TAPPA[29] | 2007 | ToPASeq R package | graphite gene-gene networks (ToPASeq)<br>user's pathways (via ToPASeq) | pathway identification | MA<br>RNASeq | p-value of DE per pathway<br>Gene-level statistics for DE of genes | Two conditions | no |

Here we propose a new method to estimate the activity within a pathway that uses biological knowledge on cell signaling to recode individual gene expression values (and/or gene mutations) into measurements that ultimately account for cell functionalities caused by the activity of the pathway. Specifically, we estimate the level of activity of stimulus-response sub-pathways (signaling circuits thereinafter) within signaling pathways, which ultimately trigger cell responses (e.g. proliferation, cell death, etc.) The activity values of these canonical circuits connected to the activation/deactivation of cell functionalities can be considered multigenic mechanistic biomarkers that can easily be related to phenotypes and provide direct clues to understand disease mechanisms and drug mechanisms of action (MoA). Therefore, we designate this method as canonical circuit activity analysis (CCAA).

## Results

### Data pre-processing

RNA-seq counts for 12 cancer types listed in Table 2 were downloaded from The Cancer Genome Atlas (TCGA) data portal (https://tcga-data.nci.nih.gov/tcga/). In order to detect possible batch effects, principal component analysis (PCA) were calculated. The samples were plotted in the PCA representation by sequencing center, plate, cancer type and project. Only a clear batch effect by sequencing center and cancer was found (Figure S1A to S1E, upper panel), that was corrected by the application of the COMBAT [21] method (Figure S1F to S1J, lower panels). Then, the 538 samples of the Kidney renal clear cell carcinoma (KIRC) dataset were further normalized using TMM [22] to account for RNA composition bias. Normalized data were used as input for the CCAA method.

**Table 2.** Cancers used in this study with the number of samples sequenced of both tumour biopsy and normal adjacent tissue.

| **TCGA Identifier** | **Cancer** | **Primary tumor** | **Normal adjacent tissue** | **Ref.** |
| --- | --- | --- | --- | --- |
| BLCA | Bladder Urothelial Carcinoma | 301 | 17 | [10] |
| BRCA | Breast invasive carcinoma | 1057 | 113 | [11] |
| COAD | Colon adenocarcinoma | 451 | 41 | [12] |
| HNSC | Head and Neck squamous cell carcinoma | 480 | 42 | [13] |
| KIRC | Kidney renal clear cell carcinoma | 526 | 72 | [14] |
| KIRP | Kidney renal papillary cell carcinoma | 222 | 32 | [15] |
| LIHC | Liver hepatocellular carcinoma | 294 | 48 | - |
| LUAD | Lung adenocarcinoma | 486 | 55 | [16] |
| LUSC | Lung squamous cell carcinoma | 428 | 45 | [17] |
| PRAD | Prostate adenocarcinoma | 379 | 52 | [18] |
| THCA | Thyroid carcinoma | 500 | 58 | [19] |
| UCEC | Uterine Corpus Endometrial Carcinoma | 516 | 23 | [20] |

### Estimation of the specificity of the CCAA method

In order to estimate the false positive rate, we generated different sets of indistinguishable samples that were randomly divided into two groups which were compared to try to find differentially activated circuits. Given that the compared groups are composed of the same type of individuals, any significant difference in sub-pathway activity found in the comparisons would be considered a false positive of the method. Real and simulated samples were used for this purpose (see Methods) and the ratio of false positives was always very low, far below the conventional alpha value of 0.05 (see Figure S2).

### Estimation of the sensitivity of the CCAA method

In order to obtain an estimation the true positive rate of the CCAA method, we compared cancer samples versus the corresponding healthy tissue in a series of contrasts with different sizes (N=50,100,200 and 400 samples; see Methods) from which we expect differences in cancer-associated pathways. Two different cancer types, KIRC and BRCA, were used to avoid biases derived from using only a specific type of cancer. We have used two definitions of cancer associated pathways, one of them taken from KEGG (composed of 14 pathways belonging to the Cancer pathways category, see Table 3), and the other one that contains 49 pathways curated by experts (Table 4). Figure S3 shows how, except in the case of very small datasets in which the statistical power of the method for detecting significant differences is limited, the proposed CCAA methodology clearly identifies significant changes for both cancers in the two cancer pathway definitions used.

**Table 3.** KEGG cancer pathways

| **KEGG identifier** | **Name** |
| --- | --- |
| hsa04010 | MAPK signaling pathway |
| hsa04310 | Wnt signaling pathway |
| hsa04350 | TGF-beta signaling pathway |
| hsa04370 | VEGF signaling pathway |
| hsa04630 | Jak-STAT signaling pathway |
| hsa04024 | cAMP signaling pathway |
| hsa04151 | PI3K-Akt signaling pathway |
| hsa04150 | mTOR signaling pathway |
| hsa04110 | Cell cycle |
| hsa04210 | Apoptosis |
| hsa04115 | p53 signaling pathway |
| hsa04510 | Focal adhesion |
| hsa04520 | Adherens junction |
| hsa03320 | PPAR signaling pathway |

**Table 4.** Curated cancer pathways

| **KEGG identifier** | **Name** |
| --- | --- |
| hsa04014 | Ras signaling pathway |
| hsa04015 | Rap1 signaling pathway |
| hsa04010 | MAPK signaling pathway |
| hsa04012 | ErbB signaling pathway |
| hsa04310 | Wnt signaling pathway |
| hsa04330 | Notch signaling pathway |
| hsa04340 | Hedgehog signaling pathway |
| hsa04350 | TGF-beta signaling pathway |
| hsa04390 | Hippo signaling pathway |
| hsa04370 | VEGF signaling pathway |
| hsa04630 | Jak-STAT signaling pathway |
| hsa04064 | NF-kappa B signaling pathway |
| hsa04668 | TNF signaling pathway |
| hsa04066 | HIF-1 signaling pathway |
| hsa04068 | FoxO signaling pathway |
| hsa04020 | Calcium signaling pathway |
| hsa04024 | cAMP signaling pathway |
| hsa04022 | cGMP-PKG signaling pathway |
| hsa04151 | PI3K-Akt signaling pathway |
| hsa04152 | AMPK signaling pathway |
| hsa04150 | mTOR signaling pathway |
| hsa04110 | Cell cycle |
| hsa04114 | Oocyte meiosis |
| hsa04210 | Apoptosis |
| hsa04115 | p53 signaling pathway |
| hsa04510 | Focal adhesion |
| hsa04520 | Adherens junction |
| hsa04530 | Tight junction |
| hsa04540 | Gap junction |
| hsa04611 | Platelet activation |
| hsa04620 | Toll-like receptor signaling pathway |
| hsa04621 | NOD-like receptor signaling pathway |
| hsa04650 | Natural killer cell mediated cytotoxicity |
| hsa04660 | T cell receptor signaling pathway |
| hsa04662 | B cell receptor signaling pathway |
| hsa04670 | Leukocyte transendothelial migration |
| hsa04062 | Chemokine signaling pathway |
| hsa04910 | Insulin signaling pathway |
| hsa04920 | Adipocytokine signaling pathway |
| hsa03320 | PPAR signaling pathway |
| hsa04912 | GnRH signaling pathway |
| hsa04915 | Estrogen signaling pathway |
| hsa04914 | Progesterone-mediated oocyte maturation |
| hsa04919 | Thyroid hormone signaling pathway |
| hsa04916 | Melanogenesis |
| hsa05200 | Pathways in cancer |
| hsa05231 | Choline metabolism in cancer |
| hsa05202 | Transcriptional misregulation in cancer |
| hsa05205 | Proteoglycans in cancer |

### Comparison to other available PAA methods

The performance of our method was compared to other PAA methods that provide different definitions of sub-pathways and distinct algorithms to calculate a score for them. From the list in (Table 1) we used eight methods that satisfy two basic conditions: they can be applied to RNA-seq data and there is software available for running them. These are: DEAP [38], subSPIA [32], using their own software, and topologyGSA [31], DEGraph [6], clipper [5], TAPPA [29], PRS [40], PWEA [30], using the implementation available in the topaseq package [41]. Figure 1 represents the true positive and true negative ratios obtained for any of the methods compared (See Methods). While most of the pathway activity definitions are reasonably specific, with true negative ratios over 95% (except clipper, topologyGSA and PWEA, probably because they define sub-pathways unconnected with cell functionality), the sensitivity is generally low (in most cases below 50%). When the curated list of cancer pathways (see Table 4) is used, the performance of some methods improves but still, the sensibility is in general low (clearly below 75%, see Figure S4).

**Figure 1.**
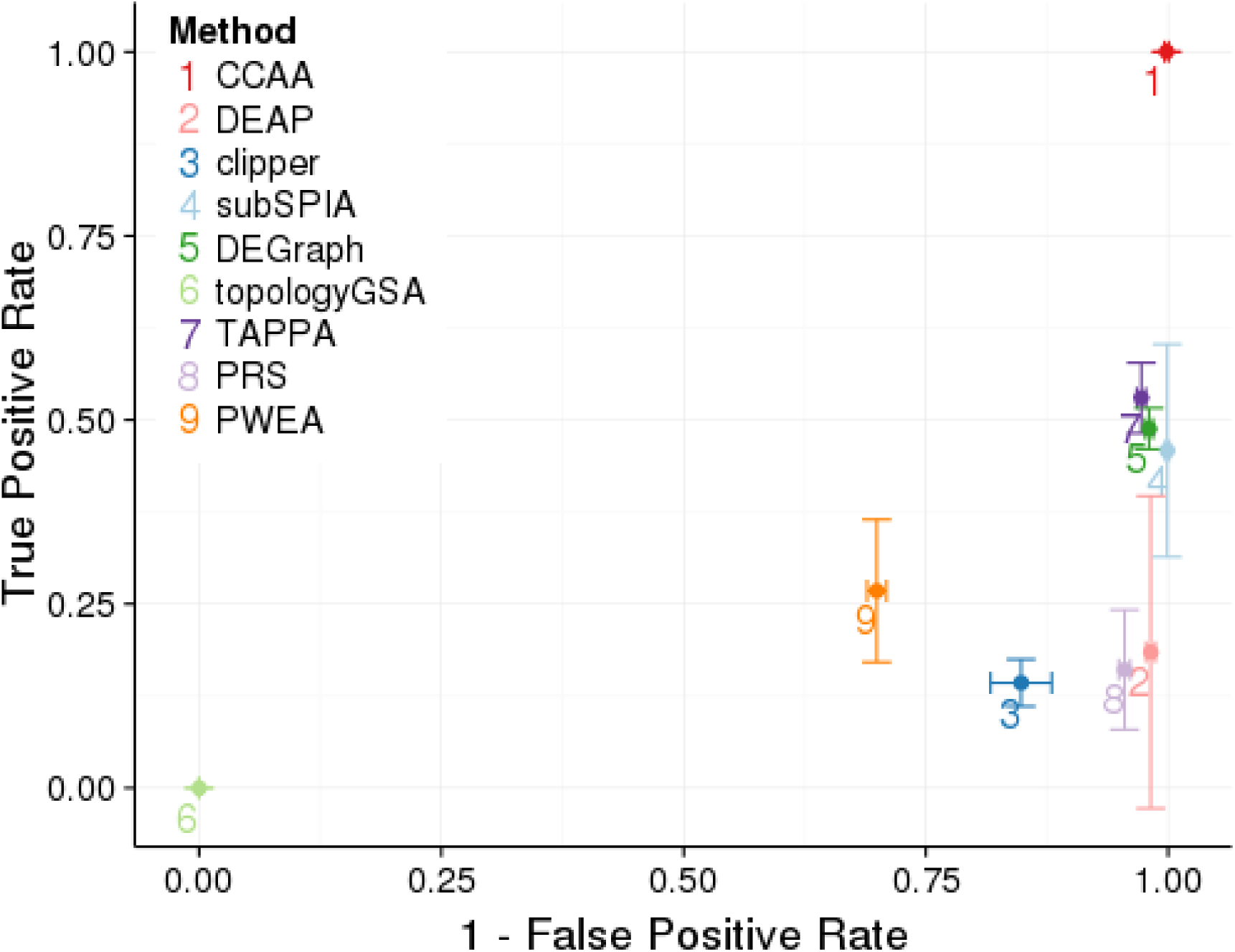
Comparison of performances of the different methods for defining pathways and calculating its activity. CCAA is compared to DEAP [38], subSPIA [32], using their own software, and topologyGSA [31], DEGraph [6], clipper [5], TAPPA [29], PRS [40], PWEA [30], using the implementation available in the topaseq package [41]. The true positive rate has been estimated averaging the proportion of significant cancer KEGG pathways (Table 3) across the 12 cancers analyzed and is represented in the Y axis. Vertical bars in each point represent 1 SD of the true positive rate for the corresponding method. The false positive rate was estimated from 100 comparisons of groups (N=25) of identical individuals, randomly sampled from each cancer. The results obtained in the 12 cancers are used to obtain a mean value and an error. The X axis represents 1-the false positive rate. Horizontal bars represent in each point represent 1 SD of the false positive rate for the corresponding method.

From the technical standpoint, the CCAA method can handle loops in the pathway topology, a feature absent in most PAA methods (see Table 1) allowing a more comprehensive description of the circuit activity.

These results demonstrates that all the PAA methods analyzed, except ours, are not properly capturing the biological signal and consequently failed to detect cancer pathway activities when cancer and normal tissues were compared, across twelve different cancer types.

### A case example with kidney renal clear cell carcinoma

To demonstrate the utility of this approach in defining the activity of canonical signaling circuits as highly reliable mechanistic biomarkers that, in addition, account for important disease outcomes such as survival, kidney renal clear cell carcinoma (KIRC) [14] data was used. In addition, survival data available on patients were used to demonstrate that the activity of many of the selected circuits is significantly related to the prognostic of the disease.

Firstly, 526 cancer samples were compared against the 72 available controls of normal kidney tissue adjacent to the primary tumors (See Table 2). The comparison was made at the level of canonical circuits (see Methods), effector circuits and functions (using both Uniprot and GO annotations). As expectable, given the large number of differentially expressed genes between the cancer and the healthy tissue [14], a large number of signaling circuits present a significant differential activation between the compared conditions (4966 with a FDR-adjusted p-value < 0.01; See Table S1).

Focusing on effector circuits, this signaling interplay is reduced to 870 significant changes in the intensity of signal reception (with a FDR-adjusted p-value < 0.01; See Table S2). These effector nodes significantly trigger 71 cell functionalities (according to Uniprot general definitions, see Table S3, which summarize 320 more detailed cell functionalities according to GO definitions, see Table S4; both with a FDR-adjusted p-value < 0.01). Figure 2 summarizes the different functions dysregulated by circuits in different KEGG cancer pathways (see Table 3) and the corresponding impact on patient’s survival. Figure S5 expands this summary to the set of curated cancer pathways listed in Table 4. Although some functionalities are quite general descriptions of cellular biological processes and others can be consequences of the extreme deregulation process occurring in cancer cells, a considerable number of them can be clearly linked to tumorigenic processes and can easily be mapped to cancer hallmarks [43].

**Figure 2.**
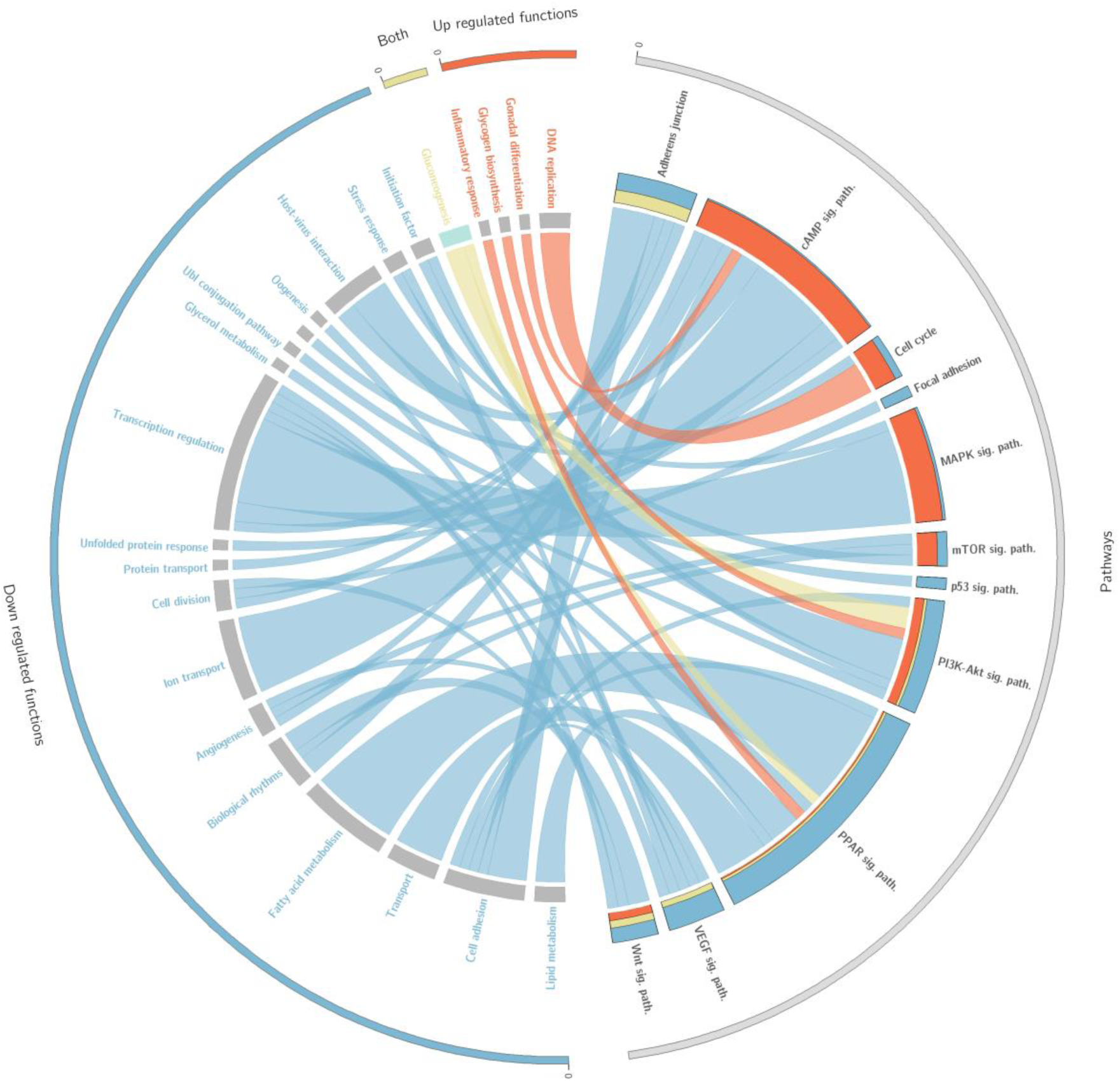
Circos plot that summarizes the relationships between effectors within pathways and the functions triggered by them. Only cancer KEGG pathways (Table 3) related to functions significantly related to survival are represented here. On the right side appear the effector circuits grouped according to the pathway they belong to. There is a histogram per pathway that represents the proportion of effector pathways upregulated (red), downregulated (blue) and dysregulated in both directions (yellow). On the left side of the circo appear the functions triggered by the effector circuits divided into those which are significant when are up-regulated (red), when are down-regulated (blue) or when both situations occur (yellow). For each function there is a band that indicates the prognostic of its deregulation, which can be good (green) or bad (grey).

### Circuits that trigger cancer hallmarks determine patient survival

Since survival data was among the clinical information available survival analysis of the significant effector circuits, and functions listed in Tables S1, S2, S3 and S4) was carried out. This analysis provides an independent validation of the involvement of several cell functionalities, as well as several signaling circuits that trigger them, in cancer pathogenesis.

Survival analysis discovered a total of 310 effector circuits whose dysregulation is significantly associated to good or poor cancer prognostic (Table S5). These circuits trigger a total of 31 general cell functionalities, according to Uniprot definitions (Table S6) that can be expanded to 108 more detailed GO definitions (Table S7), which are significantly related to patient’s survival.

The main cancer hallmark is sustained proliferation [43]. A clear example of effector circuit related to this hallmark is the *CCNA2,* from the AMPK signaling pathway, whose high levels of activity are significantly associated to bad prognostic in the patients in which triggers the *Cell division* function (Figure S6A). Actually, there is a significant increase in the activity of the *CCNA2* effector circuit as cancer stage progresses (Figure S6C). In fact, dysregulated genes were recently identified in this sub-pathway that might be potential biological markers and processes for treatment and etiology mechanism in KIRC [44]. Another similar example is the effector circuit ending in node *CDK2*, *CCNE1* from the p53 signaling pathway, and triggering the *Cell cycle* function, whose increased activity is significantly associated to bad prognostic in KIRC patients (Figure S7A and S7B). In addition, there is a significant increase in the activity of the *CDK2*, *CCNE1* effector circuit as cancer stage progresses (Figure S7C). Recently, *CDK2*, *CCNE1* genes were described as cancer prognostic factors [45]. When the association is carried out at the function level, there are two Uniprot functions (Table S6) representative of sustained proliferation hallmark: *Mitosis* (FDR-adjusted p-value 1.7x10^-12^) and *DNA replication* (FDR-adjusted p-value=5.9x10^-8^), whose upregulation is significantly associated to bad prognostic (See Figures S7A and S7B).

Another cancer hallmark is the activation of metastasis and invasion, favored when the Uniprot function *Cell adhesion* decreases. Figure S7C depicts a clear association between the downregulation of *Cell adhesion* and the poorer prognostic in patients (FDR-adjusted p-value=4.4x10^-5^).

The third classical cancer hallmark in solid tumors is the induction of angiogenesis. *Angiogenesis* appears as significantly associated to survival in both Uniprot and GO annotations (Tables S6 and S7). Figure S8D depicts a significant relationship between the upregulation of *Positive regulation of angiogenesis* and higher patient’s mortality (FDR-adjusted p-value=2.9x10^-2^). Actually, the downregulation of the opposite term, *Negative regulation of angiogenesis*, is also associated to bad prognostic, as expected, although with marginal significance (FDR-adjusted p-value=0.055).

Finally, the CCAA method also detects the well-known Warburg effect, the observed increased uptake and utilization of glucose, documented in many human tumor types [43, 46]. Our functional analysis clearly predicts a bad prognostic for reduced *gluconeogenesis* (FDR-adjusted p-value = 8.96x10^-6^, see Table S6). Actually, it has recently been suggested a novel mechanism of cancer cell death by augmenting the gluconeogenesis pathway via mTOR inhibitors [47].

In addition, the CCAA method detects several terms whose perturbed activity seem a consequence of the dedifferentiation process that occur in kidney cancer cells, such as the down-activation of *Sodium/potassium transport* (FDR-adjusted p-value=2.95x10^-9^), Sodium transport (FDR-adjusted p-value=8.96x10^-6^) and, the general term Transport (FDR-adjusted p-value= 6.52x10^-5^) (see Table S6).

### Cancer progression driven by specific circuits instead of specific genes

An additional advantage of using CCAA is that the signaling circuits that trigger the functions in this particular cancer can be easily traced back. *DNA replication* is an example of function that can easily be mapped to the *sustained proliferative signaling* cancer hallmark [43]. The increase in the activity of this function is significantly related with poor prognostic (FDR-adjusted p-value=5.94x10^-8^). Three effector circuits belonging to the *Cell cycle* and the *p53 pathways* (See Figure 3 and Table S6) are the ultimate responsible for the activation of this function. Moreover, it has been described that dysregulation of different genes within the same pathway may have a similar impact on downstream pathway function [48, 49]. Figure 4 demonstrates how the CCAA method can detect the same functional consequence (activation of DNA replication) caused by distinct, non-recurrent, differential gene expression patterns in two different cancers (BRCA and KIRC). The detection of the specific circuits and the particular gene activities involved in the tumorigenesis process has enormous therapeutic implications.

**Figure 3.**
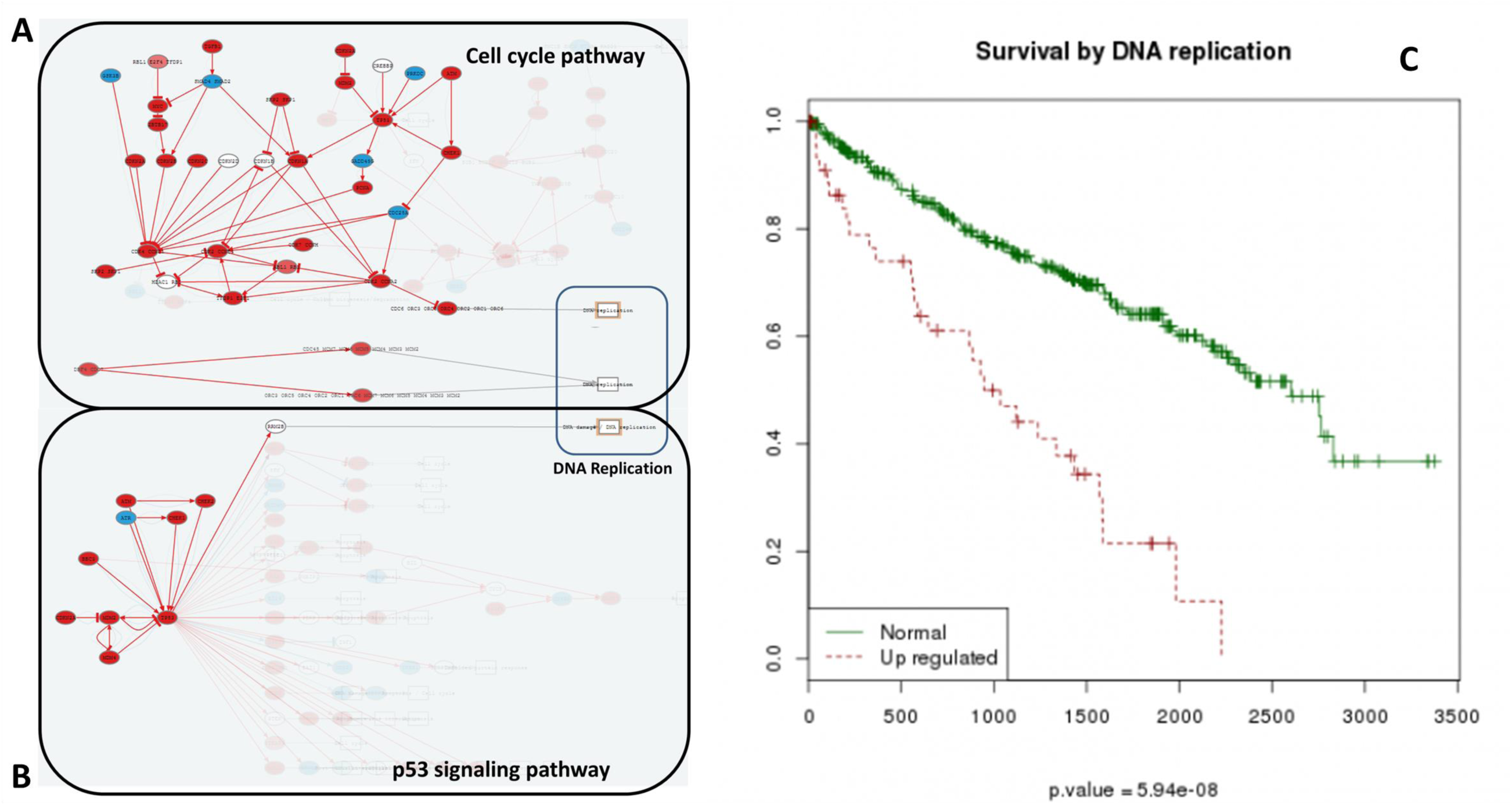
Increase of *DNA replication* is related to bad prognostic. Effector nodes in two pathways trigger *DNA replication* in KIRC, as detected by the Hipathia program (http://hipathia.babelomics.org). Genes in red represent genes upregulated in the cancer with respect to the corresponding normal tissue; genes in blue represent downregulated genes and genes with no color were not differentially expressed. A) Cell Cycle signaling pathway with three effector circuits highlighted, one of them ending in the node containing proteins *CDC6, ORC3, ORC5, ORC4, ORC2, ORC1* and *ORC6,* the second one ending in node with proteins *CDC45, MCM7, MCM6, MCM5, MCM4, MCM3* and *MCM2* and the last one ending in node with proteins *ORC3, ORC5, ORC4, ORC2, ORC1, ORC6, MCM7, MCM6, MCM5, MCM4, MCM3* and *MCM2.* B) p53 signaling pathway with the effector circuit ending in protein *RRM2B* highlighted. C) Survival Kaplan-Meier (K-M) curves obtained for Uniprot function *DNA replication*.

**Figure 4.**
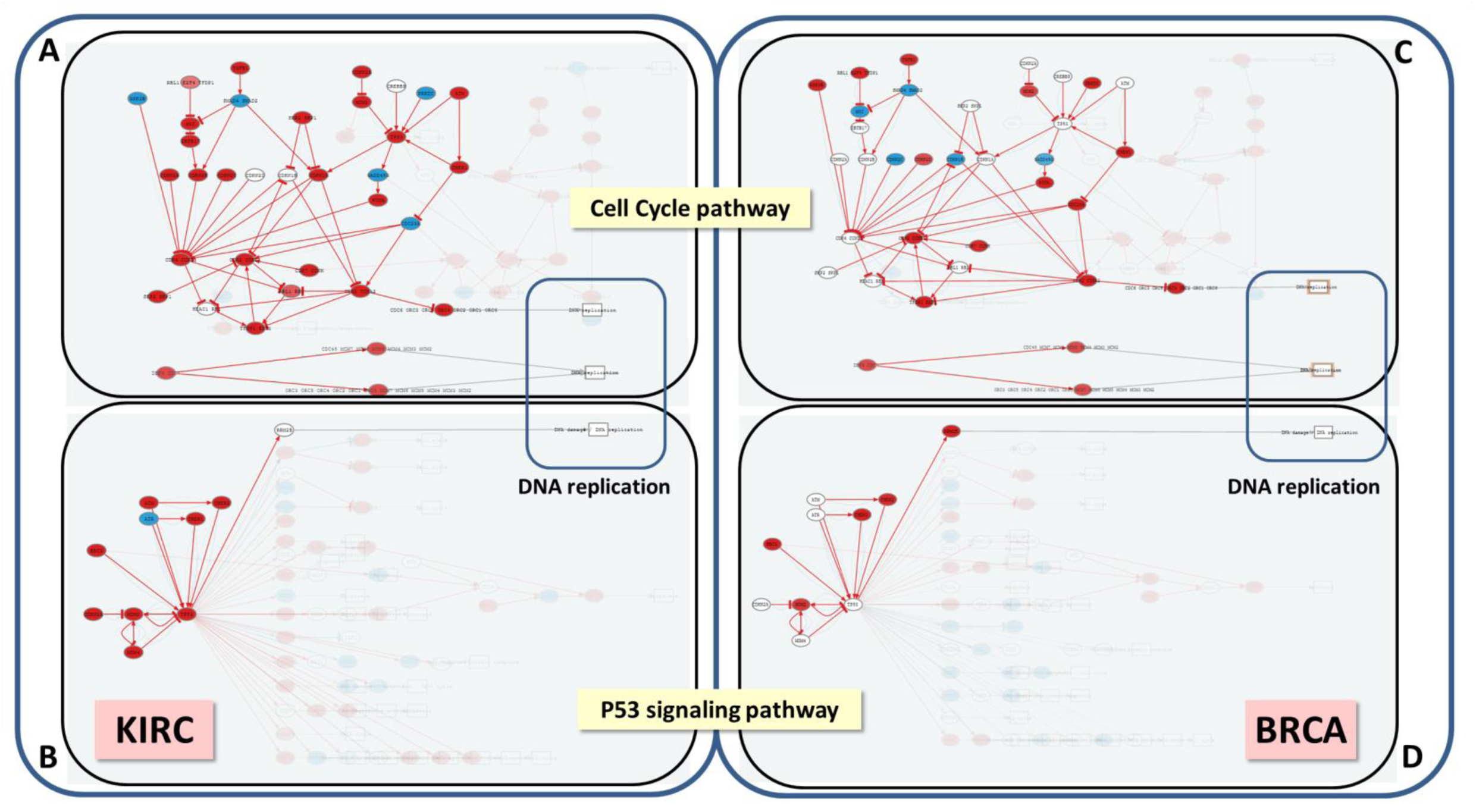
DNA replication is triggered by the same circuits in KIRC and BRCA, but using a different pattern of gene activation. The Hipathia program (http://hipathia.babelomics.org) detected a total of four effector circuits in two pathways, Cell Cycle and P53 signaling, that are used by both cancers to trigger DNA replication. Arrows in red represent activated circuits. Genes in red represent genes upregulated in the cancer with respect to the corresponding normal tissue; genes in blue represent downregulated genes and genes with no color were not differentially expressed. Squares at the end of the circuit represent the cell functions triggered by the circuits. A) Cell Cycle signaling pathway in KIRC with three effector circuits activated (highlighted), one of them ending in the node containing proteins *CDC6, ORC3, ORC5, ORC4, ORC2, ORC1* and *ORC6*, the second one ending in node with proteins *CDC45, MCM7, MCM6, MCM5, MCM4, MCM3* and *MCM2* and the last one ending in node with proteins *ORC3, ORC5, ORC4, ORC2, ORC1, ORC6, MCM7, MCM6, MCM5, MCM4, MCM3* and *MCM2*. B) P53 signaling pathway in BRCA with the effector circuit ending in protein *RRM2B* highlighted. C) Cell Cycle pathway in BRCA with the same effector circuits activated that in KIRC, but using a different set of gene activations. D) P53 signaling pathway in BRCA with the same effector circuit activated that in KIRC, but using a different set of gene activations.

## Discussion

Models of pathway activity bridge the gap between conventional approaches based on single-gene biomarkers, or functional enrichment methods, and more realistic, model-based approaches. Models use biological knowledge available on relevant biological modules (such as signaling pathways) to explain how their perturbations ultimately cause diseases or responses to treatments. Therefore, such perturbations (initially gene expression changes) can be related to disease mechanisms or drug MoAs [50, 51].

A unique feature of the CCAA method is that, if the analysis is made at the level of cell functionality, the changes in the activity detected can be traced back to the circuits in order to discover which ones are triggering the action and what genes are the ultimate causative agents of such functional activity changes. Therefore, the resulting models can be used to suggest and predict the effect of interventions (KOs, drugs or over-expressions) on specific genes in the circuits so as to find suitable clinical targets, predict side effects, speculate off-target activities, etc. Depending on the scenario studied, such interventions can be more general or more personalized.

Another relevant feature missing in the rest of PAA methods (Table 1) is the possibility of obtaining individual values of circuit, effector or function activities for each sample. This opens the door to obtaining patient-specific personalized functional profiles connected to the corresponding signaling circuits.

Since clinical data are available at the TCGA repository, we were able to find significant associations of specific pathway activities to patient survival, proving thus the validity of PAA methodology to capture cell processes involved in disease outcome.

Finally, it is worth mentioning that the integration of information on protein functionality in the model, if it is available, is straightforward. (See Methods for details). Other omic data (methylomics data, Copy Number Variation, etc.) could also be easily introduced in the model providing they could be coded as proxies of presence and/or integrity of the protein.

## Methods

### Data source and processing

We used 12 cancer types from The Cancer Genome Atlas (TCGA) data portal (https://tcga-data.nci.nih.gov/tcga/) in which RNA-seq counts for healthy control samples were available in addition to the cancer samples: Bladder Urothelial Carcinoma (BLCA) [10], Breast invasive carcinoma (BRCA) [11], Colon adenocarcinoma (COAD) [12], Head and Neck squamous cell carcinoma (HNSC) [13], Kidney renal clear cell carcinoma (KIRC) [14], Kidney renal papillary cell carcinoma (KIRP) [15], Liver hepatocellular carcinoma (LIHC), Lung adenocarcinoma (LUAD) [16], Lung squamous cell carcinoma (LUSC) [17], Prostate adenocarcinoma (PRAD) [18], Thyroid carcinoma (THCA) [19] and Uterine Corpus Endometrial Carcinoma (UCEC) [20] (Table 2).

Since TCGA cancer data has different origins and underwent different management processes, non-biological experimental variations (batch effect) associated to Genome Characterization Center (GCC) and plate ID must be removed from the RNA-seq data. The COMBAT method [21] was used for this purpose. Then, we applied the trimmed mean of M-values normalization method (TMM) method [22] for data normalization. The resulting normalized values were entered to the pathway activity analysis method.

### Modelling framework

Modelling of pathway activity requires initially of a formal description of the relationships between proteins within the pathway, which can be taken from different pathway repositories. Here KEGG pathways [23] are used, but any other repository could be used instead, as Reactome [24] or others. It also requires of a way to estimate the activation status of each protein, which accounts for the intensity of signal they can transmit along the pathway.

A total of 60 KEGG pathways (see Table 5), which include 2212 gene products that participate in 3379 nodes, are used in this modelling framework. It must be noted that any gene product can participate in more than one node (even in different pathways) and a node can contain more than one gene product. Pathways are directed networks in which nodes (composed by one or more proteins) relate to each other by edges. Only two different kinds of relation between nodes are considered: activations and inhibitions. In KEGG pathways, edges define different types of protein interactions that include phosphorilations, ubiquitinations, glycosilations, etc., but they include a label indicating if they act as activations or inhibitions.

**Table 5.** KEGG pathways modeled in this study

| **KEGG identifier** | **Name** |
| --- | --- |
| hsa04014 | Ras signaling pathway |
| hsa04015 | Rap1 signaling pathway |
| hsa04010 | MAPK signaling pathway |
| hsa04012 | ErbB signaling pathway |
| hsa04310 | Wnt signaling pathway |
| hsa04330 | Notch signaling pathway |
| hsa04340 | Hedgehog signaling pathway |
| hsa04350 | TGF-beta signaling pathway |
| hsa04390 | Hippo signaling pathway |
| hsa04370 | VEGF signaling pathway |
| hsa04630 | Jak-STAT signaling pathway |
| hsa04064 | NF-kappa B signaling pathway |
| hsa04668 | TNF signaling pathway |
| hsa04066 | HIF-1 signaling pathway |
| hsa04068 | FoxO signaling pathway |
| hsa04020 | Calcium signaling pathway |
| hsa04071 | Sphingolipid signaling pathway |
| hsa04024 | cAMP signaling pathway |
| hsa04022 | cGMP-PKG signaling pathway |
| hsa04151 | PI3K-Akt signaling pathway |
| hsa04152 | AMPK signaling pathway |
| hsa04150 | mTOR signaling pathway |
| hsa04110 | Cell cycle |
| hsa04114 | Oocyte meiosis |
| hsa04210 | Apoptosis |
| hsa04115 | p53 signaling pathway |
| hsa04510 | Focal adhesion |
| hsa04520 | Adherens junction |
| hsa04530 | Tight junction |
| hsa04540 | Gap junction |
| hsa04611 | Platelet activation |
| hsa04620 | Toll-like receptor signaling pathway |
| hsa04621 | NOD-like receptor signaling pathway |
| hsa04622 | RIG-I-like receptor signaling pathway |
| hsa04650 | Natural killer cell mediated cytotoxicity |
| hsa04660 | T cell receptor signaling pathway |
| hsa04662 | B cell receptor signaling pathway |
| hsa04664 | Fc epsilon RI signaling pathway |
| hsa04666 | Fc gamma R-mediated phagocytosis |
| hsa04670 | Leukocyte transendothelial migration |
| hsa04062 | Chemokine signaling pathway |
| hsa04910 | Insulin signaling pathway |
| hsa04922 | Glucagon signaling pathway |
| hsa04920 | Adipocytokine signaling pathway |
| hsa03320 | PPAR signaling pathway |
| hsa04912 | GnRH signaling pathway |
| hsa04915 | Estrogen signaling pathway |
| hsa04914 | Progesterone-mediated oocyte maturation |
| hsa04921 | Oxytocin signaling pathway |
| hsa04919 | Thyroid hormone signaling pathway |
| hsa04916 | Melanogenesis |
| hsa04261 | Adrenergic signaling in cardiomyocytes |
| hsa04270 | Vascular smooth muscle contraction |
| hsa04722 | Neurotrophin signaling pathway |
| hsa05200 | Pathways in cancer |
| hsa05231 | Choline metabolism in cancer |
| hsa05202 | Transcriptional misregulation in cancer |
| hsa05205 | Proteoglycans in cancer |
| hsa04971 | Gastric acid secretion |
| hsa05160 | Hepatitis C |

In order to transmit the signal along the pathway, a protein needs: first, to be present and functional, and second, to be activated by other protein. Preferably, the activity of the proteins should be inferred from (phospho)proteomic and chemoproteomic experiments [25], however, the production of these types of data still results relatively complex [26]. Instead, an extensively used approach is taking the presence of the mRNA corresponding to the protein as a proxy for the presence of the protein [5-8, 26, 27]. Therefore, the presence of the mRNAs corresponding to the proteins present in the pathway is quantified as a normalized value between 0 and 1. Second, a value of signal intensity transmitted through a protein is computed, taking into account the level of expression of the corresponding mRNA and the intensity of the signal arriving to it. The net value of signal transmitted across the pathway corresponds to the signal values transmitted by the last proteins of the pathway that ultimately trigger the cell functions activated by the pathway.

### Decomposing pathways into circuits

Pathways are represented by directed graphs, which connect input (receptor) nodes to output (effector) nodes. The signal arrives to an initial input node and is transmitted along the pathway following the direction of the interactions until it reaches an output node that triggers an action within the cell. Thus, from different input nodes the signal may follow different routes along the pathway to reach different output nodes. Within this modelling context, a canonical circuit is defined as any possible route the signal can traverse to be transmitted from a particular input to a specific output node (see Figure 5, left).

**Figure 5.**
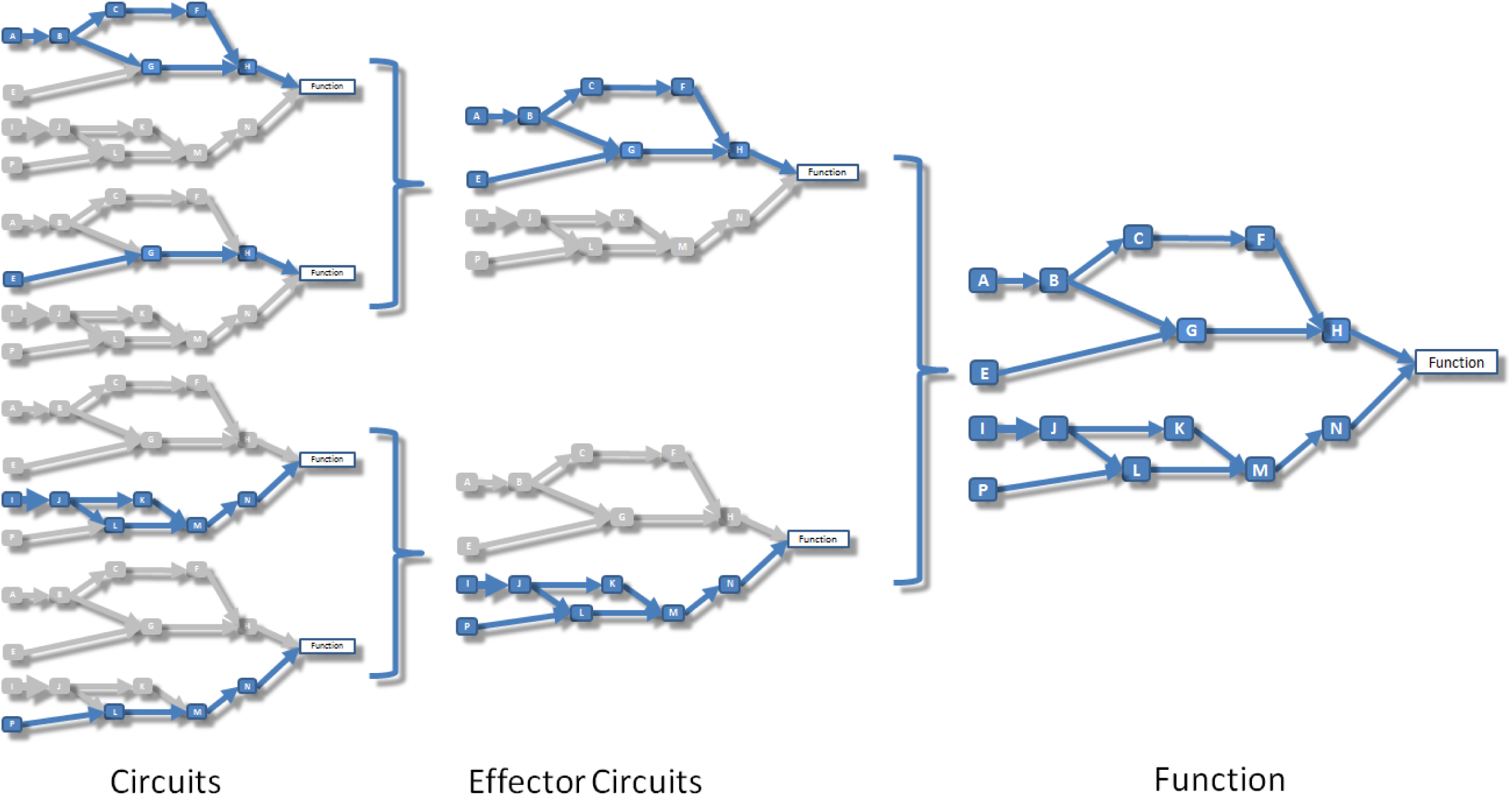
Schema that illustrates the relationship between circuits, effector circuits and functions. Left: signaling circuits, which are canonical sub-pathways that transmit signals from a unique receptor to a unique effector node. Center: effector circuits that represent the combined activity of all the signals that converge into a unique effector node. Right: functional activity that represents the combined effect of the signal received by all the effectors that trigger a particular cell function.

Output nodes at the end of canonical are the ultimate responsible to carry out the action the signal is intended to trigger in the cell. Then, from a functional viewpoint, an effector circuit can be defined as a higher-level signaling entity composed by the collection of all the canonical circuits ending in an unique output (effector) node (see Figure 5, center). When applied to effector circuits, the method returns the joint intensity of the signal arriving to the corresponding effector node.

A total of 6101 canonical circuits and 1038 effector circuits can be defined in the 60 pathways modelled.

### Computing the circuit activity

The methodology proposed uses gene expression values as proxies of protein presence values, and consequently of potential protein activation values [5-8, 26, 27]. The inferred protein activity values are then transformed into node activity values using the information on node composition taken from KEGG. KEGG defines two types of nodes: plain nodes, which may contain one or more proteins, whose value is summarized as the percentile 90 of the values of the proteins contained in it, and complex nodes, for which the minimum value of the proteins contained (the limiting component of the complex), is taken as the node activity value.

Once the node activity values have been estimated, the computation of the signal intensity across the different circuits of the pathways is performed by means of an iterative algorithm beginning in the input nodes of each circuit. In order to initialize the circuit signal we assume an incoming signal value of 1 in the input nodes of any circuit. Then, for each node n of the network, the signal value is propagated along the nodes according to the following rule:

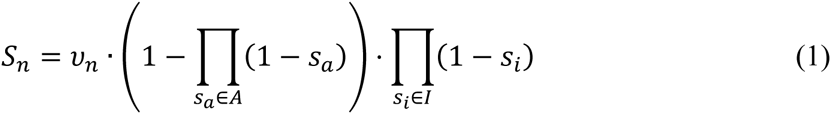

where *A* is the total number of signals arriving to the node from activation edges, *I* is the total number of signals arriving to the node from inhibition edges, and *v_n_* is the normalized value of the current node *n*.

The algorithm to compute the transmission of the signal along the network is a recursive method based on the Dijkstra algorithm [28]. Each time the signal value across a node is updated in a recursion and the difference with the previous value is greater than a threshold, all the nodes to which an edge arrives from the current updated node are marked to be updated. The recursion continues until the update in the values is below the threshold. The advantage if using an iterative method is that the signal becomes steady even in cases of loops in the pathway topology, allowing a more precise estimation of circuit activities. Many PAA methods simply cannot handle with loops and artificially disconnect them or even remove them from the calculations [5, 6, 8, 29-32]. Figure 6 represents the computation of the intensity of signal transmission across a node, and exemplifies in a simple scenario how the signal is transmitted across a circuit.

**Figure 6.**
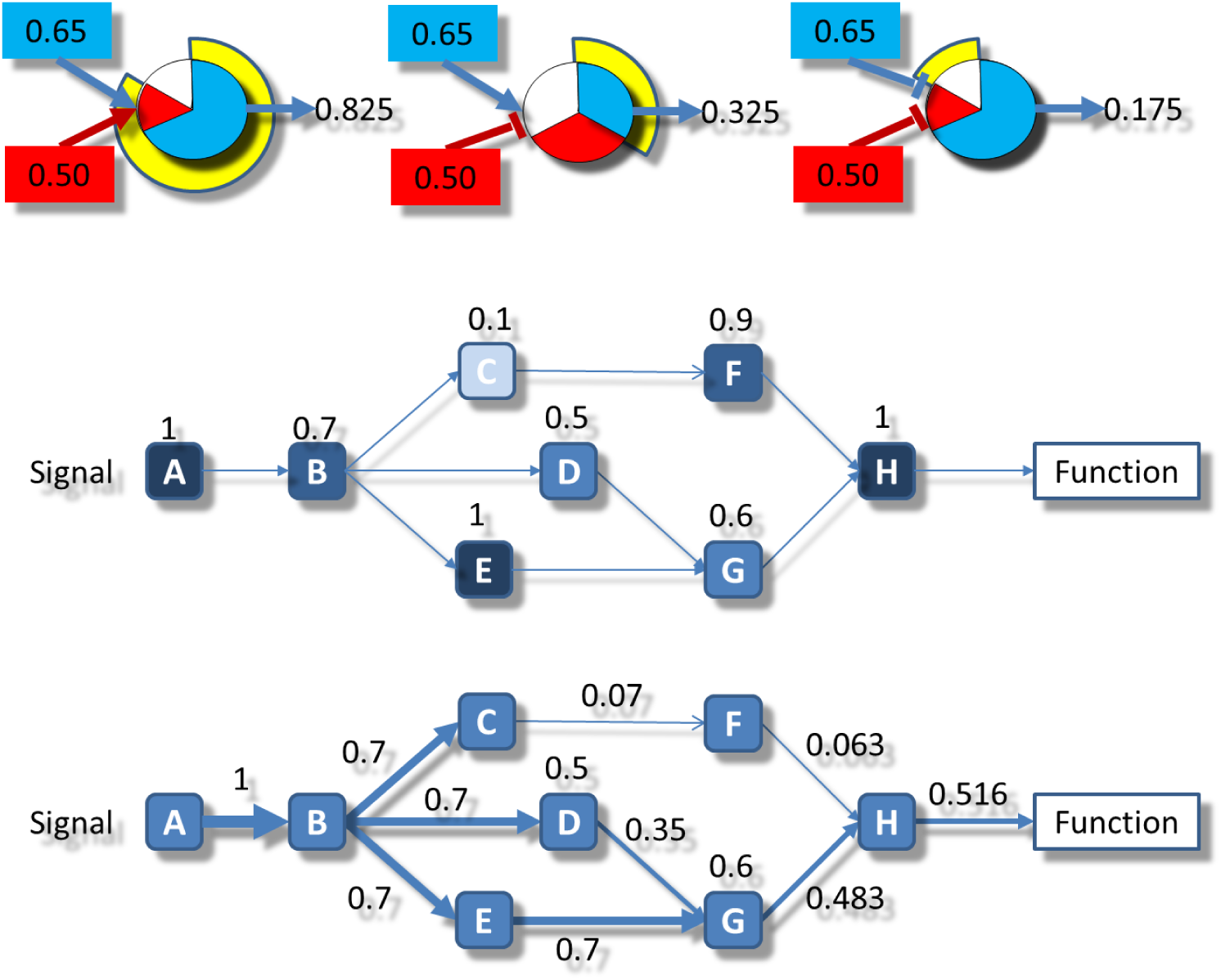
Schematic representation of the signal propagation algorithm used. Upper part: the three types of activity transmitted: left) the combination of two activations, center) the combination of an activation and an inhibition and right) the combination of two inhibitions. Central part: the normalized values of gene expression are assigned to the corresponding nodes in the circuits. Lower part: the signal starts with a value of 1 in the receptor node A and is propagated by multiplying the weights assigned to each node in the central part following the rules depicted in the upper part.

### Effector circuits and functional analysis

Effector nodes at the end of the circuits trigger specific functions in the cell. These functions are defined here based on the annotations of the proteins contained in the effector node. Gene Ontology [33] (GO) terms corresponding to the biological process ontology (February 16, 2016 release) and molecular function keywords of Uniprot [34] (release of September 21, 2015) are used.

The signal intensity received by the effector node can be propagated to the functions triggered by them following the same rationale of signal propagation along the circuits. Figure 5 illustrates how effector circuits are composed by different canonical circuits and how functions can be triggered by several effector circuits.

### Straightforward integration of transcriptomic and genomic data

Finally, the integration of genomic and transcriptomic data in the proposed modeling framework of signaling pathways is straightforward. In order to transmit the signal a protein needs to be present (gene expressed) and to be functional (harboring no impairing mutations). Genomic data can be integrated with transcriptomic data to infer combined gene activity and integrity (and consequently potential functionality). In the simplest approach [9] the normalized expression value of genes harboring mutations is multiplied by 0 if the pathogenicity (e.g. SIFT [35], PolyPhen [36]) and conservation indexes (e.g. phastCons [37]) are beyond a given threshold (taking into account the inheritance mode), or if the consequence type of the mutation (stop gain, stop loss, and splicing disrupting) is deleterious *per se*, because it is considered to produce a non-functional protein. The HiPathia program enables the analysis of mutations found in standard variant files (VCF) from whole exome/genome sequencing experiments in combination with gene expression values.

### Specificity of the method of canonical circuit activity analysis (CCAA)

To estimate the false positive rate, different groups of N identical individuals were generated and further divided into two datasets that were compared to each other for finding differentially activated circuits. This comparison was repeated 2000 times for different data sizes (N = 20, 50, 100, 200 and 400 individuals) in three different scenarios: i) N individuals were randomly sampled among KIRC patients; ii) For each gene *g*, an empirical distribution of gene expression values was derived from the patients of the KIRC dataset, with mean µ_g_ and variance σ^2^_g_. Then, N individuals were generated by simulating their gene expression values as random numbers sampled from a normal distribution N(µ_g_,σ^2^_g_); iii) N individuals were generated by simulating their gene expression values as random numbers from a normal distribution N(0.5, 0.05).

Since the individuals involved in the comparison were taken either from the same type of samples or were generated in the same way, any differential activation found can be considered a false positive. The comparisons were carried out for both, circuits and effector proteins.

### Sensitivity of the Canonical Circuit Activity Analysis (CCAA) method

To estimate the true positive rate, we tested a scenario in which biological differences are expected. For this purpose, we used the two 2 cancers in Table 2 with more individuals, BRCA [11] and KIRC [14]. For each of the two cancers we generated 100 datasets of N=50,100,200 and 400 samples by sampling randomly both the normal and tumor samples in such a way that the normal/tumor proportion remained the same as in the original dataset (Table 2). In total, we generated 2x100x4 = 800 datasets. CCAA was calculated at the level of signaling circuits and effector circuits for both datasets. The true positive rate was estimated as the number of cancer pathways containing one or more differentially activated circuits divided by the total number of cancer pathways. Although a gold standard is always difficult in this type of scenario, we can expect changes in the 14 cancer pathways, as defined in KEGG (Cancer pathways category, see Table 3). Additionally, we produced an extended table of 49 cancer pathways curated by expert collaborators from the Valencia Institute of Oncology (IVO) (Table 4).

### Comparison with other available methods for defining and scoring pathway activity

We compared the reliability of the CCAA method proposed here to other proposals for defining sub-pathways and for calculating an activity score for them. Among the methods listed in Table 1 only nine could be applied to RNA-seq data and have software available for running them. These are: DEAP [38], subSPIA [32], using their own software, and SPIA [39], topologyGSA [31], DEGraph [6], clipper [5], TAPPA [29], PRS [40], PWEA [30], implemented in the topaseq package [41]. The relative performance of the methods compared was derived from the estimation of their ratios of false positives and false negatives in a similar way than above. In order to estimate the false positives rate 12 cancer datasets (Table 2) were used. For each cancer, 50 patients were randomly sampled 100 times. Any sampled set is divided into two equally sized subsets that are subsequently compared. Then, the 100 values obtained for each cancer are used to determine a mean value and a SD for the false positives ratio. The same 12 cancers (Table 2) were used to estimate the true positive rates. For each cancer versus normal tissue comparison the number of significant cancer pathways was calculated and divided by the total number of cancer pathways. The ratios were calculated for both the 14 cancer pathways as defined in KEGG (Cancer pathways category, see Table 3) and the extended list of 49 curated cancer pathways (Table 4).

### Survival in cancer

The KIRC TCGA samples contain survival information among the clinical data available. Kaplan-Meier (K-M) curves [42] were estimated using the function *survdiff* from the *survival* R package (https://cran.r-project.org/web/packages/survival/) for each signaling circuit, each effector circuit and each cell function (either Uniprot or GO definitions) with a significant difference of activity when cancers were compared to the corresponding controls. Specifically, the 10% of individuals presenting the highest (or lowest) activity were compared to the rest of them.

### Availability of data and materials

A user-friendly web server that runs the code for carrying out the CCAA method is freely available at http://hipathia.babelomics.org.

The R code implementing the method is available at https://github.com/babelomics/hipathia.

## Acknowledgements

We are very indebted to Drs. José Costa, from the Yale University School of Medicine, and Jose Antonio Lopez Guerrero, from the Valencian Institute of Oncology (IVO), for their valuable comments and help in the biological interpretation of the results found.

## Author contributions

M.R.H. and J.C.C. developed the method and analyzed data; A.A. and C.C. analyzed data; F.S. and J.C.C. developed analytical tools; and J.D. conceived the method and wrote the paper.

## Grant support

This work is supported by grants BIO2014-57291-R from the Spanish Ministry of Economy and Competitiveness and “Plataforma de Recursos Biomoleculares y Bioinformáticos” PT13/0001/0007 from the ISCIII, both co-funded with European Regional Development Funds (ERDF); PROMETEOII/2014/025 from the Generalitat Valenciana (GVA-FEDER); Fundació la Marató TV3 (ref. 20133134); and EU H2020-INFRADEV-1-2015-1 ELIXIR-EXCELERATE (ref. 676559) and EU FP7-People ITN

Marie Curie Project (ref 316861)

## Additional files

**Additional File 1: Figure S1**. PCA plots of the samples to discover batch effects. **Figure S2**. False positive ratio of the CCAA method proposed, obtained as the proportion of signaling circuits that present significant differential activity when identical datasets are compared. **Figure S3**. True positive ratio of CCAA method proposed obtained as the proportion of cancer pathways with one or more signaling circuits with a significant differential activity found by comparing cancer cases to their corresponding normal tissue samples, for which real differences are expected. **Figure S4**. Comparison of performances of the different methods for defining pathways and calculating its activity. **Figure S5**. Circos plot that summarises the relationships between effectors within pathways and the functions triggered by them. **Figure S6.** Example of effector circuit significantly associated to bad prognostic in KIRC. **Figure S7**. Example of effector circuit significantly associated to bad prognostic in KIRC. **Figure S8**. Survival Kaplan-Meier (K-M) curves obtained for Uniprot and GO functions.

**Additional File 2. Table S1**. Canonical circuits differentially activated between cancer and the normal tissue. **Table S2**. Effector circuits differentially activated between cancer and the normal tissue. **Table S3**. Unitprot functions differentially activated between cancer and the normal tissue. **Table S4**. Gene Ontology functions differentially activated between cancer and the normal tissue. **Table S5**. Effector circuits associated to patient survival. **Table S6**. Uniprot functions associated to patient survival. **Table S7**. Gene Ontology functions associated to patient survival.

